# A self-assembled protein nanoparticle serving as a one-shot vaccine carrier

**DOI:** 10.1101/2020.09.16.299149

**Authors:** Ten-Tsao Wong, Gunn-Guang Liou, Ming-Chung Kan

## Abstract

In this paper, we are exploring the role of an amphipathic helical peptide in mediating the self-assembly of a fusion protein into a protein nanoparticle and the application of the nanoparticle as a one-shot vaccine carrier. Out of several candidates, an amphipathic helical peptide derived from M2 protein of type A influenza virus is found to stimulate high antigenicity when fused to a fluorescent protein genetically. This fusion protein was found to form protein nanoparticle spontaneously when expressed and purified protein stimulates long-lasting antibody responses in single immunization. Through modeling peptide structure and nanoparticle assembly, we have improved this vaccine carrier in complex stability. The revised vaccine carrier is able to stimulate constant antibody titer to a heterologous antigen for at least six months in single immunization. The immune response against a heterologous antigen can be boosted further by additional immunization in spite of high immune responses to carrier protein.

## Introduction

Subunit vaccine is a safe alternative to traditional inactivated or attenuated vaccines, but its efficacy is often hindered by the low antigenicity of recombinant protein. Different approaches are utilized to resolve this issue, among them, virus like particle (VLP) and self-assembled protein nanoparticle (SAPN) are considered the best platforms for subunit vaccine development(*1*). VLP is assembled from a recombinant capsid protein alone without the genomic nucleic acid and bound nucleocapsid protein so it is non-infectious(*2*). The size of VLP is ranged between 20-200 nm that facilitates both draining efficiently to lymph node and also uptake by antigen presenting cells like dendritic cell and macrophage(*3*). The other benefit of VLP based vaccine is the induction of B cell receptor clustering when presenting repetitive antigen to B cell, a function that can activate antibody class-switch and somatic hypermutation in a T cell dependent mechanism(*4*). Not only the virus like particle can be used for vaccine directly, heterologous antigen can be presented in the particle surface through genetic fusion(*5*). The Universal flu vaccine candidate, ectopic M2 domain (M2e), was genetically fused with Hepatitis B core antigen(HBc) and assembled into a nanoparticle that provides full protection to homologous flu strain(*6*). But the application of HBc based VLP in human vaccine development is restricted by the pre-existing anti-HBc antibody present in the 450 million chronic HBV carriers and the population exposed to HBV infection(*7*). An artificially designed SAPN may avoid the effect of the existing antibody. A SAPN assembled from protein constituted by two coiled-coil domains that form trimer and pentamer respectively can be assembled into nanoparticles with specific sizes(*8, 9*). This SAPN stimulates strong immune responses to target protein fused to the terminal of constituting monomer after 3 immunizations even without adjuvant, but this immunity waned gradually (*10*).

The green fluorescent protein is a member of fluorescent protein family that are structurally conserved and emit fluorescent light from a chromophore when excited by photons of shorter wavelength(*11*). The shared features of fluorescent proteins including a sturdy barrel shaped structure constituted by 11 β-sheets and an enclosed chromophore that emits fluorescent light when excited (*12*). The function of the barrel shell is to provide a well organized chemical environment to ensure the maturation of chromophore and protects it from hostile elements(*13*). So it is conceivable that the protein sequences among fluorescent protein family members in the barrel shell are highly variable and fluorescent proteins possess desirable biophysical properties can be selected using directed evolution(*11, 14–17*). The applications of fluorescent protein have been expanded into multiple areas beyond live imaging which includes serving as biological sensors (*18, 19*), or detectors for protein-protein interaction or protein folding(*20, 21*).

Amphipathic α-helical peptide (AHP) forms hydrophilic and hydrophobic faces when folded and is often identified in proteins related to phospholipid membrane interaction. The N-terminal amphipathic helical peptide is required for membrane anchorage of Hepatitis C virus NS3 protein and the protease function of the NS3/NS4a complex (*22, 23*). Several anti-microbial peptides also possess amphipathic properties and function by forming membrane pores or causing membrane disruption(*24*). The amphipathic α-helical of type A influenza virus M2 protein is required for M2 protein anchorage and induces membrane curvature required for virus budding (*25, 26*).

Life protection from diseases through vaccination is considered a holy grail only achievable by some live attenuated viral vaccines with multiple doses up to date (*27*). Here in this report, we are describing the identification of a protein nanoparticle based on an amphipathic α-helical peptide (AHP) from M2 protein of type A influenza strain H5N1 and a green fluorescent protein. We have predicted the protein nanoparticle structure according to transmissive electronic microscope images using protein modeling and generated AHP mutants that provide higher stability and antigenicity to the protein nanoparticle. This modified protein nanoparticle is able to simulate long constant antibody response to an inserted heterologous antigen in single immunization without adjuvant. And this antibody response can be further boosted by additional immunization. The identification and application of this nanoparticle as a vaccine carrier was explored in this study.

## Results

### Identification of an AHP-GFP protein complex with high antigenicity and stability

As described in our patent application filed in 2015, we have tested the immunogenicity of fusion proteins composed of an AHP and a GFP(*28*). The results showed an increase of anti-GFP IgG titer ranged between 2~3 log under a two immunizations regime (Figure S1). One of the peptides, AH3, that derived from M2 protein of type A influenza strain H5N1 gives extended stability to the GFP fusion protein when compared to another peptide, AH1 (Figure S2) as well as other peptides in our study (data not shown). Since a stable protein is essential for vaccine carrier and may profits worldwide vaccination effort, we were interested in the mechanism of AH3-GFP stability and antigenicity. To study the potential mechanisms that contribute to the above mentioned properties of AH3-GFP fusion protein, we first checked the composition of AH3-GFP protein post expression and purification. One clue that led us to study the composition of AH3-GFP fusion protein is the difficulties encountered during protein purification. Unlike other fusion proteins studied, both AH3-GFP and AH5-GFP fusion proteins are mostly expressed as insoluble inclusion body and the remaining soluble protein did not bind to Ni-NTA resin under normal condition of 300mM NaCl. The AH3-GFP and AH5-GFP fusion proteins only started to bind to Ni-NTA resin after lowering the NaCl concentration from 300mM to 50mM, an indication that hydrophobic interaction may induce a protein complex with N-terminal His tag hindered from binding to Ni-NTA ligand. Also, the resistance of AH3-GFP fusion protein to hydrolysis suggested the linker between AH3 peptide and GFP is kept in a water tight complex. From these two clues, it was hypothesized that AH3-GFP fusion protein form stable protein complex through hydrophobic interaction mediated by N-terminal AH3 peptide.

### Characterization of AH3-GFP protein complex

To test the hypothesis that AH3-GFP or AH5-GFP fusion protein forms a protein complex, we first used the protein concentration tube with different molecular weight cut off (MWCO) to determine the protein complex sizes. As shown in figure 1A, when GFP protein with a molecular weight of 27kDa is able to pass through membranes with MWCO of 100kDa, 300kDa and 1000kDa freely, AH3-GFP fusion protein purified from bacterial lysate was prevented from passing through the membrane with an MWCO of 1000kDa. With a molecular weight of 33kDa, the purified AH3-GFP protein need to form a complex with more than 30 monomers to be excluded from passing a membrane with a 1000kDa MWCO. To explore further the geometric composition of the AH3-GFP protein complex, we examined the fusion protein under transmissive electronic microscope (TEM). The TEM results showed the AH3-GFP fusion protein forms a cylinder-like structure with length up to ~60 nm and a diameter around 10 nm (Fig. 1B). The difference in length suggests that the particle may be assembled along the long axis. When scanning along the long axis of the AH3-GFP particle, there is a repetitive pattern of two-one-two-one of white dots with two less visible dots on each side of the single dot. The predicted structure according to TEM images is shown in Fig.1D. We also examined protein geometric composition of AH5-GFP under TEM, but there is no clear evidence of forming higher order protein complex, suggesting AH5-GFP protein complex is not as stable as AH3-GFP to withstand the conditions during negative staining. To find the correlation between protein complex formation and antigenicity, we immunized mice with purified AH3-GFP fusion protein and the recombinant GFP protein that pass through the membrane freely. Proteins were prepared from LPS synthesis defective E. coli strain, ClearColi BL21(DE3), to avoid the interference of LPS contamination, a known TLR4 ligand. The mice were immunized with purified proteins by single intramuscular injection and sera were collected at day 7, 14, 30 and 182 to evaluate anti-GFP IgG titer by ELISA. Deoxycholate was added to test if deoxycholate in the concentration of 0.2% affects AH3-GFP antigenicity and related experiment was terminated at 30 days post immunization when it showed no effect on antigenicity of either GFP or AH3-GFP. These results suggest GFP alone is a poor antigen and only gained high antigenicity after fused with AH3 peptide (Fig. 1C).

**Figure 1.**
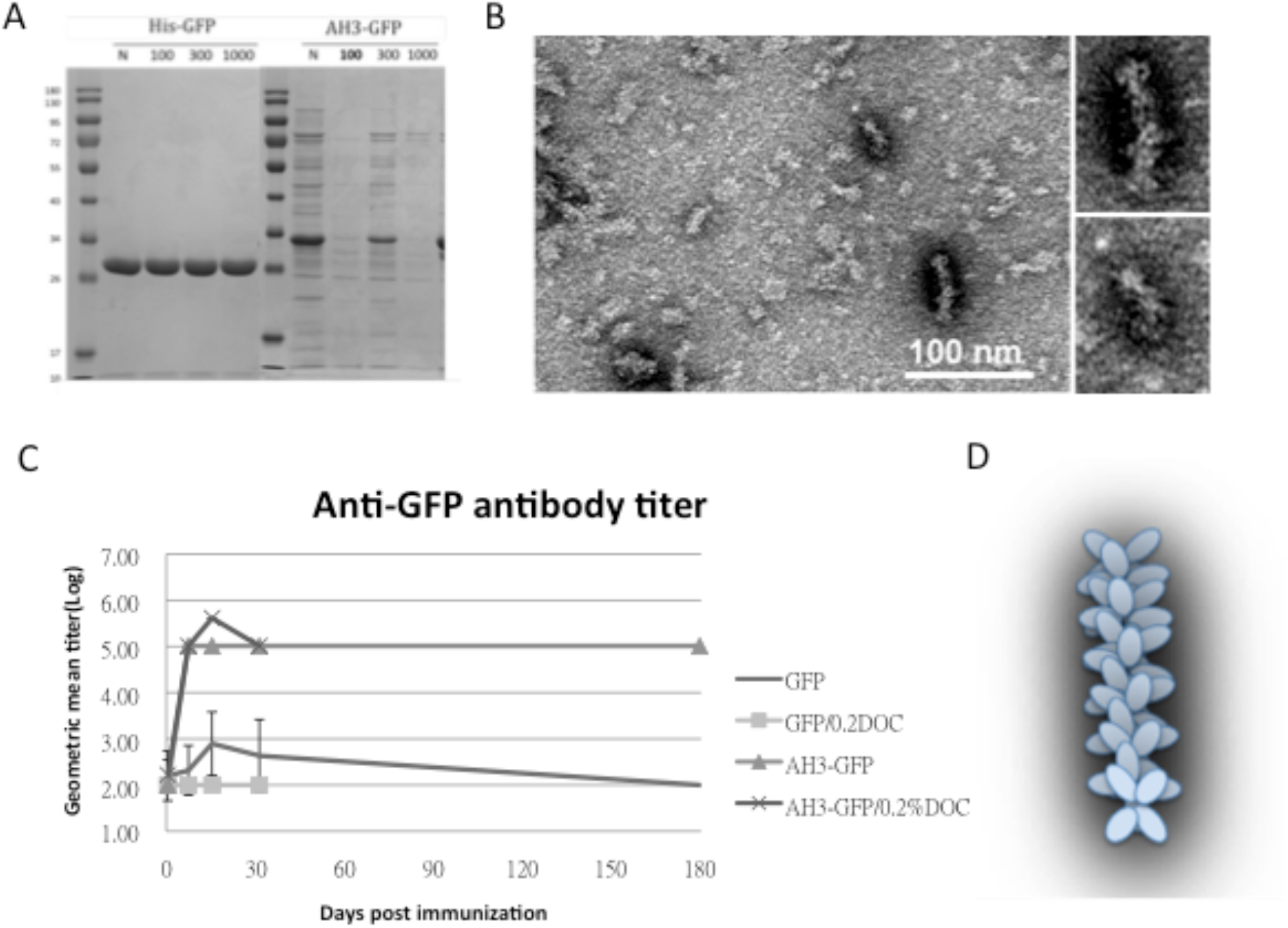
Characterization of GFP fusion protein oligomerization. (A) GFP and AH3-GFP fusion proteins purified and reconstituted in PBS were centrifuged through size exclusion membrane with different molecular weight cut off (MWCO). The filtrate was then analyzed by 12% SDS-PAGE and Coomassie blue staining (arrows are needed to point out the bands). (B) Immunogenicity of GFP and AH3-GFP fusion protein is evaluated by immunizing mice in a single muscular immunization. The anti-GFP IgG titers were followed for 6 months by ELISA. (N=5). (C) The oligomerization of AH3-GFP protein was analyzed by negative staining using a transmissive electronic microscope(TEM). Scale bar represents 100nm. Two of the representative particles are enlarged and shown in the right panels. (D) AH3-GFP protein oligomer model.

### Prediction of the AH3-GFP protein complex structure using protein modeling

To understand the potential molecular mechanism leading to the assembly of AH3-GFP nanoparticle, we tried to build a protein model based on three observations: first, the particle was assembled through hydrophobic interaction and second, the particle assembled along the long axis and the third: AH3-GFP protein particle has a repetitive three-two pattern when observed under TEM. We first assembled the two AH3 peptide as anti-parallel helices with hydrophobic sidechains of F4, F5, I8, L12 and L16 mediating intermolecular contacts. The assembled AH3 dimer created a hydrophobic core between two helices and the helix dimer is surrounded by hydrophilic sidechains from multiple lysine and arginine except one exposed hydrophobic patch, as predicted using Deepview and marked by white mesh (Fig. 2A). This hydrophobic patch can be seen only to cover one face of the dimer (Fig. 2B). When the water accessible surface of AH3 dimer was calculated, two cavities could be seen located within the hydrophobic patch that provides contact points for two Arg9 sidechains extruding from the opposite face of AH3 dimer. A second AH3 dimer can make close contact with the first dimer after turning counter clockwise looking down the hydrophobic patch for 36° and forms a tetramer (Fig. 2C). The intermolecular energy between two dimers from this model was calculated to has a ΔG of −101 kcal/mol (Fig. 2C). After adding GFP protein structures onto the AH3 tetramer model, the AH3-GFP fusion protein tetramer will form a cross-shaped assembling unit and the stacking of every AH3-GFP tetramer on top of another tetramer will extend the particle length by 2.8 nm and turning the cross by 72°. Since the GFP protein barrel diameter is ranged between 2.7~3.5 nm, the out extending GFP from AH3-GFP tetrameric cross can spatially fit with the model (Fig. 2D). Under this model, the protein nanoparticle will be extended continuously with a hydrophobic patch presenting on one end of the assembled particle constitutively and serving as a point for polymerization.

**Figure 2.**
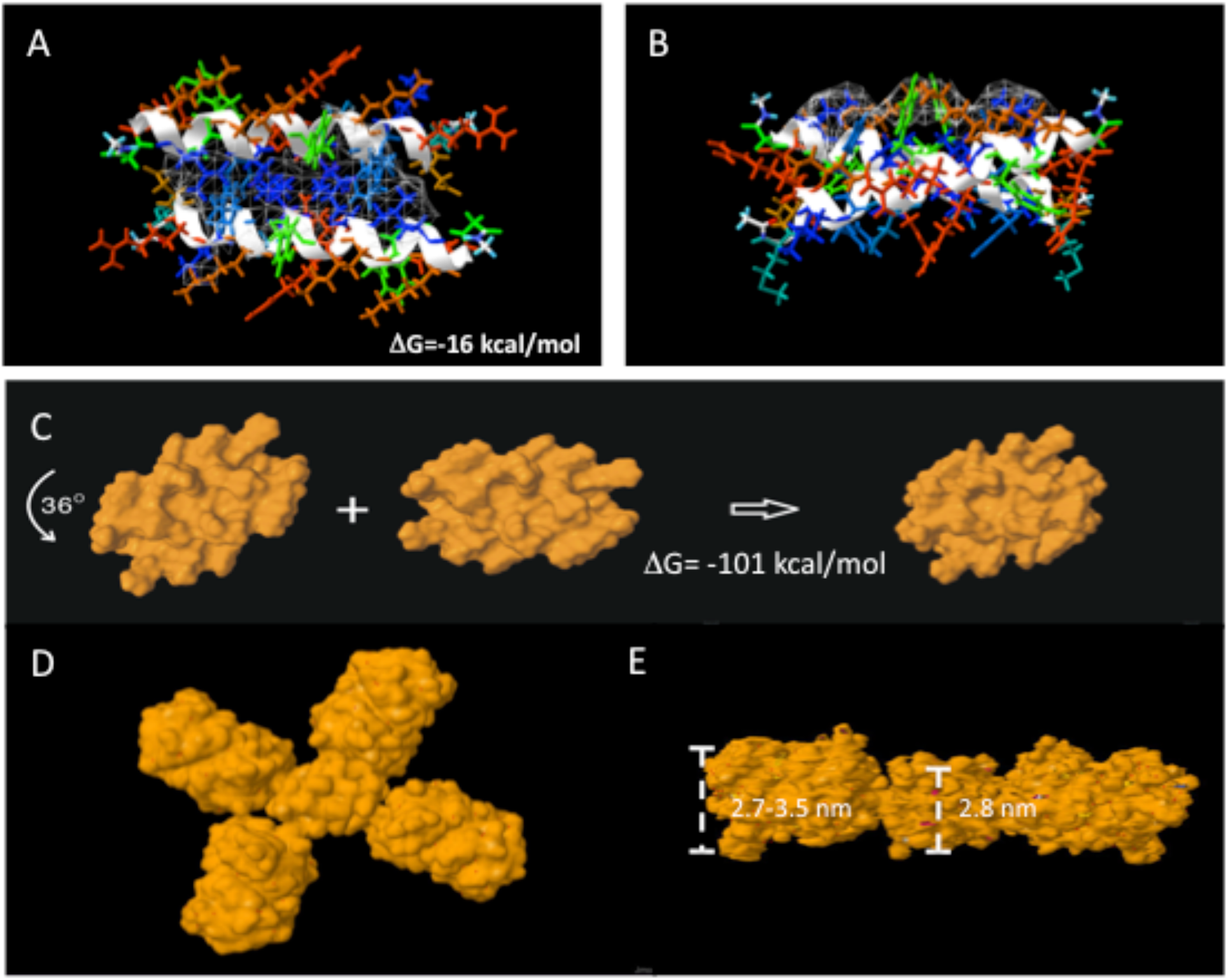
AH3-GFP protein oligomer structural modeling. (A) AH3 peptide monomer was modeled as an α-helical structure and forms anti-parallel dimer centered on hydrophobic core. The sidechain is colored according to hydrophobicity, with most hydrophobic amino acid sidechain in blue and most hydrophilic sidechain in red and spans the spectrum according to hydrophobicity. The hydrophobic patch is covered by white mesh. (B) Bottom view of AH3 dimer that shows the location of the hydrophobic patch. (C) Water accessible surface of AH3 dimer protein model and the stacking of 2^nd^ dimer onto the first one after turning counter clockwise for 36°. (D)The AH3 tetramer with 4 linked GFP molecules. (E) The AH3-GFP tetramer with two GFP molecules removed for a clear view. The distance bars showing the relative scale of molecules thickness.

### Crafting a vaccine carrier that enables heterologous antigen insertion and high stability

After proving that the AH3-GFP protein complex possesses high antigenicity, we decided to explore the application of AH3-GFP protein nanoparticle as a vaccine carrier. For the Hepatitis B core antigen, the amino acid 144 served as an insertion site for heterologous antigen fusion (*29*). GFP protein has a thermal stable structure that constituted by 11 β-strands and 3 α-helices and some of the loops between strands have been explored as insertion sites for heterologous protein for various purposes (*18, 19, 30*). Among those candidates, loop173 linking strand 8 and strand 9 was chosen because it has a high capacity for foreign peptide insertion (Fig 3A) (*30*). The original AH3-GFP recombinant protein is constructed in pET28a vector with AH3 coding region inserted C-terminal to His-tag and thrombin cleavage site followed immediately by GFP cloned from pEGFP-C2. This expression vector was low in soluble protein productivity and unable to express soluble recombinant protein when the peptide is inserted between D173 and G174. To resolve the expression and folding issues, we designed a new expression vector. First, we cloned AH3 peptide into the very N-terminal following methionine in the pET27 vector, and then to its C-terminal we inserted a synthesized sfGFP gene (*31*) with a created antigen insertion site following 175S of sfGFP. The antigen insertion site also included an 8xHis tag for recombinant protein purification. To verify vaccine carrier function, we inserted two copies of broad spectrum flu vaccine candidate, human M2 ectopic peptide (hM2e) separated by a 6 amino acid linker (Fig. 3B). The newly constructed vector was proven to be efficient for expressing soluble AH3-sfGFP-2hM2e fusion protein as a protein complex (data not shown). The AH3-sfGFP-2xhM2e protein complex under TEM is not as stable as AH3-GFP unless first cross-linking the protein preparation with a heterobifunctional protein crosslinker, sulfo-SMCC (Fig. 3C). Following the previous established AH3-GFP protein model, we were seeking strategies to create a more stable AH3-sfGFP protein complex. First, we found the mutation of Isoleucine 8 to Leucine increase the intermolecular interaction(ΔG) from −16 kcal/mol to −37 kcal/mol (Fig.3D). Second, we mutated Lysine 13 to Glutamic acid and generated additional electrostatic interactions between side chains of Glu13 with Arg10 and Arg11 (Fig. 3E).

**Figure 3.**
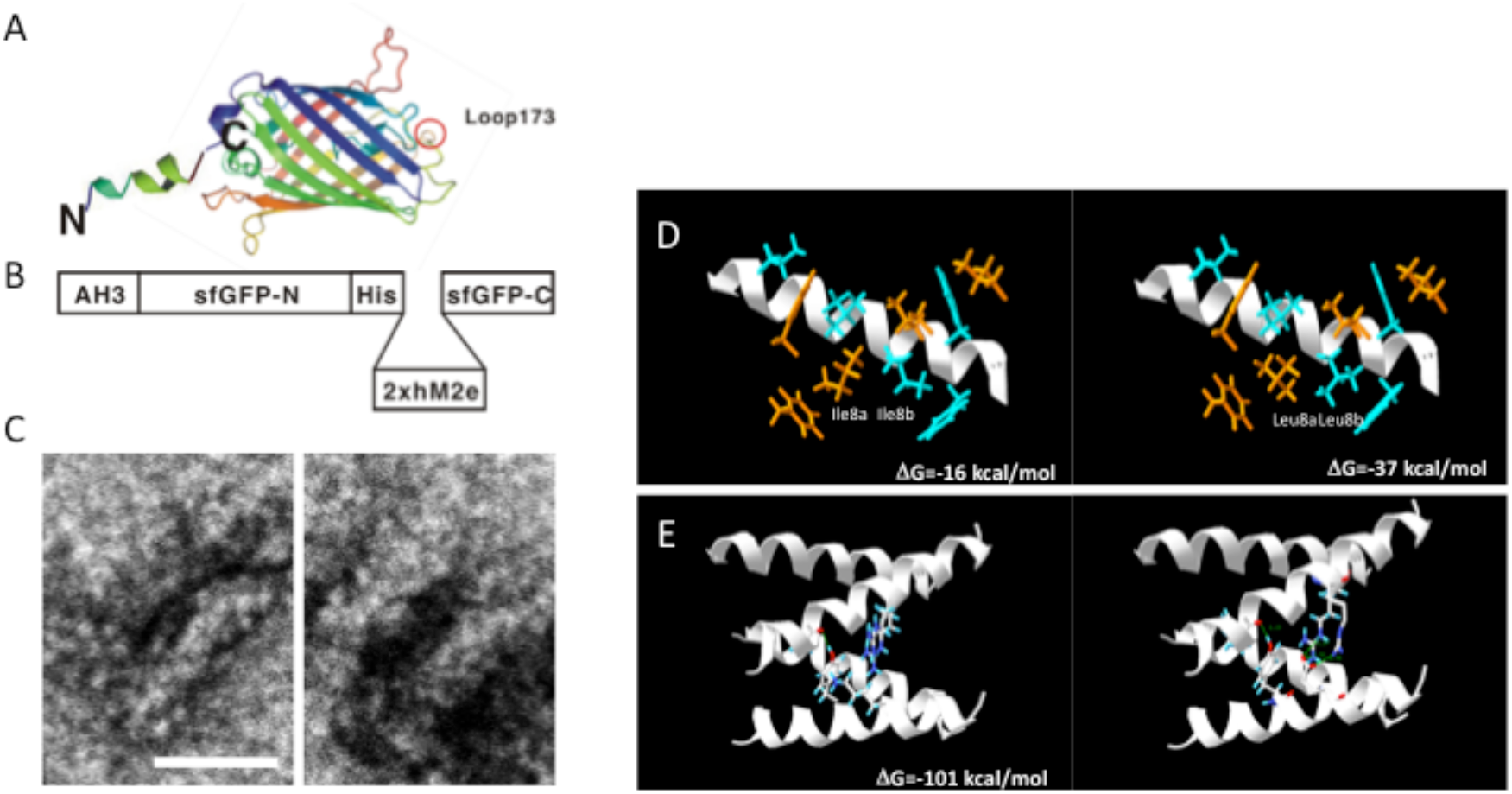
Construction of AH3-sfGFP-2xhM2e fusion protein and protein model guided AH3 mutagenesis for variants with a higher oligomer stability. (A) Graphic presentation of AH3-sfGFP fusion protein and the insertion site (loop173) marked by a circle. The 19 amino acid AH3 peptide was fused to the N-terminal of sfGFP to mediate oligomerization and a foreign antigen insertion site placed in loop173 with an 8xHis tag for purification. (B) The graphic presentation of AH3-sfGFP-2xhM2e fusion protein. (C) Analysis of AH3-sfGFP-2xhM2e protein oligomerization using TEM after crosslinking with sulfo-SMCC. The scale bar represents 20nm. (D) Comparison of AH3 and AH3 I8L (LY) mutation in dimer formation. (E) Comparison of AH3 dimers and AH3 K13E mutation dimers in forming tetramer.

To verify whether the protein modeling results are correct, including the presence of hydrophobic patch on AH3 peptide complex and a higher stability of AH3 based protein complex with I8L and/or K13E mutations. We first generated mutations in AH3 peptide in the context of AH3-sfGFP-2xhM2e construct (Fig. 4A). Our hypothesis is that the hydrophobic patch of the AH3-GFP complex will bind bacterial membrane and co-sediment with it during ultracentrifugation. And then the methodology was verified by first centrifuge bacterial lysate prepared from ClearColi culture in a centrifuge tube preloaded with 15%(w/v), 45%(w/v) and 85%(w/v) sucrose solution in the volume of 7ml, 2ml, 1ml respectively (Fig. 4B left panel). The distribution of bacterial membrane was marked by a lysochromic dye, Sudan III. The control sample contained Sudan III with lysis buffer alone (Fig. 4B lane 1). After ultracentrifugation, the bacterial membrane is sedimented to the junction of 15%/45% sucrose solution (Fig.4B lane 2). Using the same protocol, AH3-GFP was found to co-sedimented with the bacterial membrane (Fig. S3A) as well as the AH3-sfGFP-hM2e fusion protein but not a free GFP protein (Fig S3B) when the tubes were illuminated by 450nm wavelength LED light. After the protocol was verified, then the bacterial lysates of all six fusion proteins were prepared and analyzed under the same protocol. From the distribution of fluorescent protein between different percentages of sucrose solution, we have made several observations: 1. AH3-sfGFP-2xhM2e protein complex binds to bacterial membrane and cosedimented to 15%/45% junction, so as AH3 variants LY and LYRLLK (Fig. 4C lane 1). 2. Compared to lane 1, 2 and 3, when K13 is mutated to glutamic acid, this mutation decreases the interaction between protein complex and bacterial membrane (junction of 15%/45%), lane 4, 5 and 6. The protein complex of AH3 variant LYRRLE and RRLE even sedimented further to 45%/85% junction (Fig. 4C lane 5 and 6). 3. Arginine 11 is important for stabilizing AH3-K13E fluorescent protein complex, since its mutation to leucine (Fig. 4C lane 4) destabilized protein complex as shown by the increase of fluorescent in 15% sucrose fraction (Fig. 4E), compare LYRLLE vs LYRRLE and RRLE. These results confirmed the presence of hydrophobic patch in AH3-FP protein complex as predicted from our protein modeling also the role of R11 in stabilizing AH3-FP protein complex as predicted by protein modeling. What surprised us is the loss of hydrophobic patch in AH3-FP protein complex when Lysine 13 was mutated to Glutamic acid (Aspartic acid as well, data not shown) as demonstrated by the decrease of fluorescent protein distribution to 15%/45% junction. These results are consistent with our protein structure modeling that Arg11 serves as a main contact point for dimer stacking also it mediates the electrostatic interaction with mutated Glu13 (Fig. 3E).

**Figure 4.**
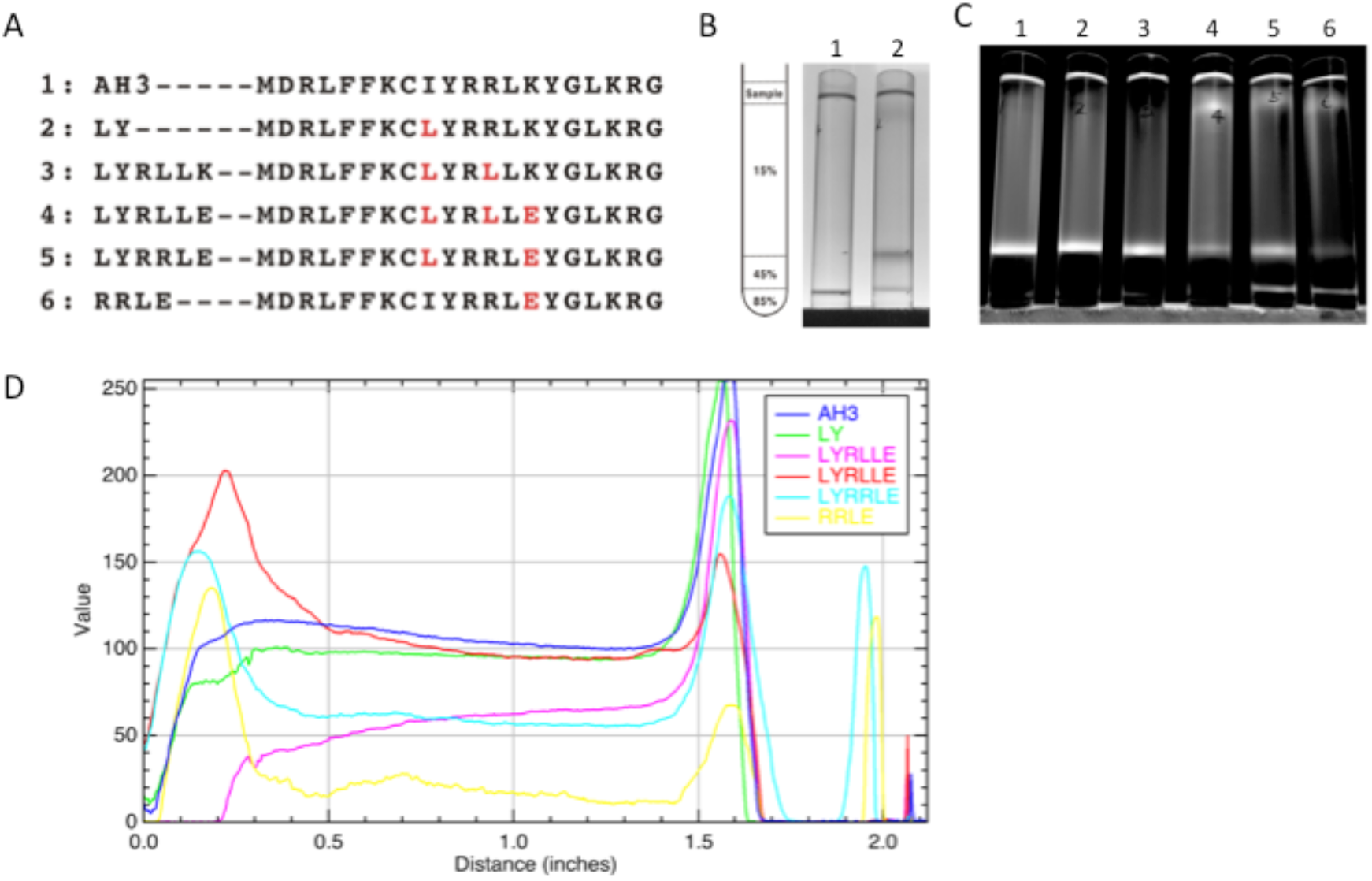
The analysis of stability and hydrophobicity of AH3-sfGFP-2xhM2e variants by sucrose density gradient. (A) List of AH3 peptide variants generated on AH3-sfGFP-2xhM2e fusion construct for sucrose density centrifuge analysis. (B) The right panel shows the graphic presentation of the centrifuge tube loaded with step sucrose solutions. The left panel shows the membrane distribution post ultracentrifugation as indicated by Sudan III staining. Lane 1 is no lysate control, lane 2 is topped with 1ml bacterial lysate. Both samples were mixed with stock staining solution by 1:100. (C) Distribution of sfGFP fusion proteins post ultracentrifugation as illuminated by 450nm LED light. (D) Quantification of fluorescent intensity along the centrifuge tube from top to bottom.

As shown in the sucrose step gradient result, the presence of hydrophobic patch enables nonspecific interaction of the AH3-GFP protein complex with phospholipid membrane. Which may restrict the free moving of protein nanoparticle and keep it from reaching draining lymph node for stimulating immunity (*4*). To compare the antigenicity of protein complexes derived from either AH3-sfGFP-2xhM2e or LYRRLE-sfGFP-2xhM2e, we immunized mice with a single injection of either recombinant proteins. Post immunization, sera were collected at day 15, 50, 90 and 202 to evaluate anti-hM2e IgG titer by ELISA. The geometric mean titer of anti-hM2e IgG reached the highest point for the AH3-sfGFP −2xhM2e group and then declined afterward. But of the LYRRLE-sfGFP-2xhM2e group, the GMT reached highest point at day 50 and remained steady up to day 90 (Fig. S4). When the individual mouse serum result is observed separately, only one out of 5 mice from AH3-sfGFP-2xhM2e group has higher anti-hM2e IgG titer at day 202 than day 15. But there are 4 out of 5 mice from the LYRRLE-sfGFP-2xhM2e group shows a higher antibody titer in day 202 compared to day 15 (Fig. 5A). These results suggest that the two point mutations of AH3 in I8L and K13E enable the formation of a stable, high antigenic protein complex that stimulates long lasting immune responses in a single immunization.

**Figure 5.**
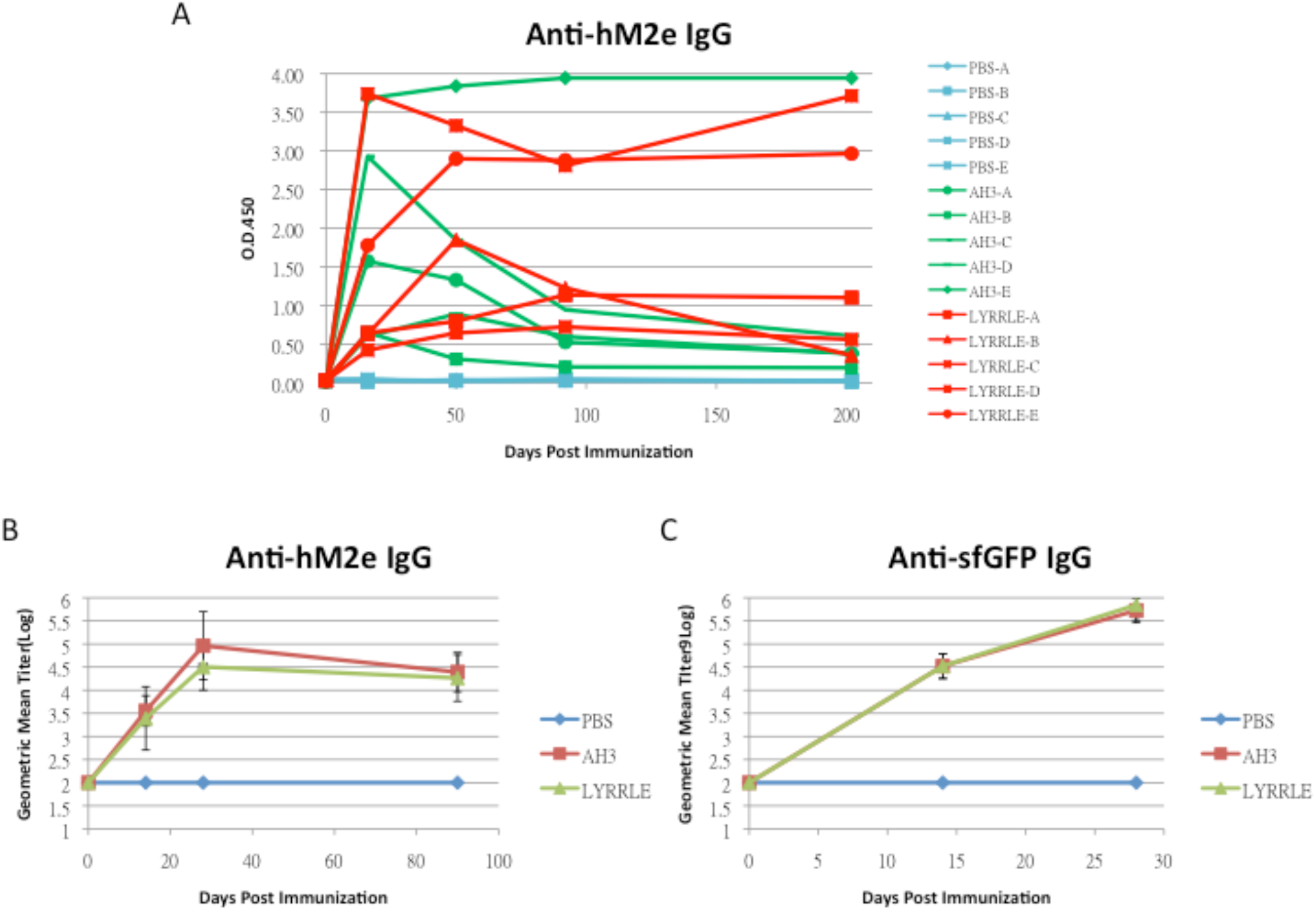
Immunization of mice with AH3-sfGFP-2xhM2e and LYRRLE-sfGFP-2xhM2e and detection of anti-hM2e IgG and anti-sfGFP IgG. (A) The mice were immunized once at day 0 by 20 μg of purified fusion proteins or PBS and then sera were collected at day 0, 15, 49, 90 and 202 for analysis to detect total anti-hM2e IgG by ELISA. Anti-hM2e total IgG titers are presented as optical density at OD450 using immune serum from each mouse diluted 1:100. (B) The anti-hM2e total IgG of mice immunized twice with 14 days apart in a prime-boost regime were followed for 3 months and analyzed by ELISA (N=5). (C) The anti-sfGFP total IgG of mice immunized by AH3-sfGFP-2xhM2e or LYRRLE-sfGFP-2xhM2e twice were evaluated 14 days post each immunization by ELISA (N=5).

Vaccine carrier like HBc based virus like particle (VLP) is often failed at boosting humoral immune responses after prime dose in a multiple doses protocol due to antigen competition. Although GFP is a protein of low antigenicity, the fusion with AH3 strongly enhances its antigenicity as shown in Figure 1C. To test if sfGFP backbone competes with inserted hM2e peptide for immune machinery, we immunized mice in a prime-boost protocol using the same protein preparations. The two consecutive injections were carried out 14 days apart and sera collected at day 14, 28 and 90 were subject to ELISA assay using either hM2e peptide or sfGFP protein as coating antigen. The result shows the IgG titer against hM2e elevated continuously after consecutive immunizations for both proteins as well as anti-sfGFP IgG titer. The result suggests that although carrier protein AH3-sfGFP also has high antigenicity, it did not interfere with the immune response against the heterologous protein, hM2e (Fig. 5B, 5C)

## Discussion

Vaccine as a tool to prevent infectious disease is the most cost effective strategy. Especially for attenuated viral vaccines like vaccinia, MMR or oral polio vaccine, they produce long lasting even life time protective immune responses but these attenuated viral vaccines took decades for development. Apparently, this strategy will not be able to timely develop a vaccine to ward off emerging global pandemic like Covid-19. Although the new vaccine technology like DNA vaccine, mRNA vaccine or adenovirus based vaccine that can quickly develop a subunit vaccine after the genomic information of pathogen become available, but the immune responses generated are often declined to base level within a year (*32–35*). This short lived immune response may expose vaccinated people to the risk of antibody dependent enhancement (ADE) that is known to be devastating and leads to vaccine failure (*11, 36*). In this study, we have created a self-assembled protein nanoparticle composed of AH3-FP and a more stable variant that stimulates long lasting antibody responses. The nature of this long lasting immune responses is not known but it may be mediated by long lasting plasma cell generated during AH3-FP immunization, the same mechanism that accounts for the lifelong protection of attenuated viral vaccines (*27*).

### Fluorescent proteins as vaccine carriers

Fluorescent protein family is a group of proteins with conserved barrel shaped structure with chromophore buried inside. Since the function of this beta strand constituted barrel is to provide a suitable environment for chromophore maturation, the primary sequence of fluorescent protein barrel is prone to mutagenesis through either direct selection (*11*) or evolution (*12*). Also, fluorescent proteins of desired biophysical properties like thermal stability or folding efficiency can be obtained through direct selection. The variability and flexibility of fluorescent protein make it a powerful tool as a vaccine carrier. First of all, the thermal stability of fluorescent protein makes AH3-FP based vaccine tolerant to high temperature, a critical property in the global vaccination initiative. Second, the AH3-FP nanoparticle platform may be expanded into several vaccine carriers that share a common format but each with distinct serum types. Third, the fast folding nature of fluorescent protein makes it a better platform to accommodate heterologous proteins. The insertion of a foreign protein in the middle of a protein structure often lead to protein destabilization and a reduction in productivity. In this aspect, AH3-FP format is suitable for the commercialization of a nanoparticle based subunit vaccine.

### Amphipathic helical peptide

In this study, the amphipathic helical peptide from M2 protein (M2AH) of type A influenza strain H5N1 is found to induce nanoparticle formation when fused with a fluorescent protein. Previous studies about M2AH focus on its role in virus budding by generating membrane curvature through embedding hydrophobic face in membrane bilayer (*26*) and anchoring M2 ectopic domain on the viral envelope as a proton pump during infection (*37*). Here we discover a new function of M2AH, serving as a nucleating center for protein complex assembly. This function is restricted to conditions when a specific M2AH is fused with fast folding proteins like fluorescent proteins. Fusing amphipathic helical peptide to other slow folding proteins tested causes misfolding of fusion protein apparently due to the aggregation induced by AH3 peptide. So, the fast folding ability of fluorescent protein is key to the AH3-FP protein nanoparticle formation. In this study, we go further to identify one point mutation in AH3 peptide, K13E, that is key to prohibit non-specific interaction of AH3-FP protein complex with cellular membrane and it plays significant role in extending the lifetime of antibody responses.

## References

1. J. Lopez-Sagaseta, E. Malito, R. Rappuoli, M. J. Bottomley, Comput Struct Biotechnol J 14, 58 (2016).

2. V. Manolova et al., European Journal of Immunology 38, 1404 (2008).

3. F. Ahsan, I. P. Rivas, M. A. Khan, A. I. Torres Suárez, Journal of Controlled Release 79, 29 (2002).

4. M. F. Bachmann, G. T. Jennings, Nature Reviews Immunology 10, 787 (2010).

5. Y. Lu, W. Chan, B. Y. Ko, C. C. VanLang, J. R. Swartz, Proc Natl Acad Sci U S A 112, 12360 (2015).

6. N. V. Ravin et al., Vaccine 33, 3392 (2015).

7. D. C. Whitacre, B. O. Lee, D. R. Milich, Expert review of vaccines 8, 1565 (2009).

8. S. Raman, G. Machaidze, A. Lustig, U. Aebi, P. Burkhard, Nanomedicine: Nanotechnology, Biology and Medicine 2, 95 (2006).

9. P. Burkhard, C. P. Karch, Nanomedicine (Lond) 12, 1529 (2017).

10. S. A. Kaba et al., Journal of immunology (Baltimore, Md.: 1950) 183, 7268 (2009).

11. R. Y. Tsien, Annual Review of Biochemistry 67, 509 (1998).

12. S. G. Dove, O. Hoegh-­-Guldberg, S. Ranganathan, Coral Reefs 19, 197 (2001).

13. C. W. Cody, D. C. Prasher, W. M. Westler, F. G. Prendergast, W. W. Ward, Biochemistry 32, 1212 (1993).

14. D. W. Close et al., Proteins 83, 1225 (2014).

15. C. Kiss, J. Temirov, L. Chasteen, G. S. Waldo, A. R. Bradbury, Protein Eng Des Sel 22, 313 (2009).

16. B. P. Cormack, R. H. Valdivia, S. Falkow, Gene 173, 33 (1996).

17. A. Crameri, E. A. Whitehorn, E. Tate, W. P. C. Stemmer, Nature biotechnology 14, 315 (1996).

18. T. V. Pavoor, Y. K. Cho, E. V. Shusta, Proc Natl Acad Sci U S A 106, 11895 (2009).

19. R. Wang et al., Appl Environ Microbiol 80, 4126 (2014).

20. S. p. Cabantous, T. C. Terwilliger, G. S. Waldo, Nature biotechnology 23, 102 (2005).

21. M. G. Romei, S. G. Boxer, Annual review of biophysics 48, 19 (2019).

22. Y. He et al., Virology 422, 214 (2012).

23. S. M. Horner, H. S. Park, M. Gale, Jr., J Virol 86, 3112 (Mar, 2012).

24. A. Tossi, L. Sandri, A. Giangaspero, Peptide Science 55, 4 (2000).

25. S. Rossman, X. Jing, G. P. Leser, R. A. Lamb, Cell 142, 902 (2010).

26. K. L. Roberts, G. P. Leser, C. Ma, R. A. Lamb, Journal of virology 87, 9973 (2013).

27. I. J. Amanna, N. E. Carlson, M. K. Slifka, N Engl J Med 357, 1903 (2007).

28. M. Kan. US20180161425A1 (2018).

29. G. P. Borisova et al., FEBS Letters 259, 121 (1989).

30. T. Kobayashi et al., PLoS One 3, e3822 (2008).

31. J.-­-D. Pédelacq, S. Cabantous, T. Tran, T. C. Terwilliger, G. S. Waldo, Nature Biotechnology 24, 79 (2006).

32. K. Modjarrad et al., The Lancet. Infectious diseases 19, 1013 (2019).

33. R. A. Feldman et al., Vaccine 37, 3326 (2019).

34. J. X. Li et al., Lancet Glob Health 5, e324 (2016).

35. F. C. Zhu et al., Lancet 389, 621 (2016).

36. H. W. Kim et al., American Journal of Epidemiology 89, 422 (1969).

37. B. Hu, S. Siche, L. Möller, M. Veit, Journal of virology 94, e01605 (2020).

