## Supplementary material for "A self-assembled protein nanoparticle serving as a one-shot vaccine carrier"

**Figure Legend**

**Figure S1.**

Anti-GFP antibody titers of mice immunized by fusion proteins composed of amphipathic helical peptide and GFP. Mice were immunized twice with 14 days apart by intramuscular injection of 20 μg of fusion protein indicated. Sera collected at day 14 to day 16 post booster immunization were used for ELISA analysis. Data were compiled from 3 different individual experiments under the same experimental procedure. The ELISA plates were coated with 10μg/ml GFP and the 2^nd^ antibody was used at a concentration of 1:2000 dilution. The other procedures were executed as standard protocol. (N=4 or 5) The result suggested the fusion of amphipathic helical peptide onto GFP increases the antigenicity of GFP by 2~3 log.

**Figure S2.**

Examining the protein stability of AH1-GFP and AH3-GFP fusion protein in ambient temperature. AH1-GFP and AH3-GFP transformed into BL21(DE3) or ClearColi competent cells for protein expression were induced at 15^o^C overnight and then following the standard protocol for protein purification. Proteins were dialyzed into 1xPBS in a dialysis membrane with MWCO of 3000 Da overnight. (A) Purified proteins were kept in 37^o^C for one week or one month and then analyzed by SDS-PAGE and Coomassie blue staining. (B) Fusion proteins from ClearColi were kept in 37^o^C one month or RT (25^o^C) for one week or 5 months and then analyzed by SDS-PAGE and Coomassie blue staining. Results from figure S2A indicates AH1-GFP fusion protein stability in 37^o^C is reduced in the presence of LPS, but not AH3-GFP. Also, from Figure S2B, the result indicates the AH3-GFP is stable in both 25 ^o^C and 37^o^C up to 5 months. But for AH1-GFP fusion protein, more than 70% of the protein was degraded into GFP in a week and completely converted to GFP in 5 months in 25 ^o^C. The results suggest AH3-GFP is stable in both ambient temperatures tested.

Figure S3.

Using a sucrose step density gradient to analyze the hydrophobicity and stability of the AH3-GFP fusion protein complex. Protein samples were loaded on top of sucrose step gradients and then centrifuged at 35K rpm for 2 hrs in a SW41 Ti rotor. After centrifugation, fluorescent protein distribution was detected under 450nm LED light and pictures taken for analysis by ImageJ from NIH. (A) The centrifuge tube was first loaded with 1ml 85% sucrose (w/v), 2 ml 45% sucrose (w/v) and 7 ml 15% sucrose (w/v) consecutively and then topped with bacterial lysate containing AH3-GFP fusion protein for analysis. (B) The centrifuge tube was loaded first with 1ml 45% (w/v) sucrose solution and then followed with 9ml of 15% (w/v) sucrose solution. Samples of 1 ml each were loaded on top of the centrifuge tube before ultracentrifugation as described. Results of two samples were shown in figure S3B: GFP and AH3-sfGFP-2xhM2e. The results showed the GFP protein alone remained in the top of the centrifuge tube, but AH3-sfGFP-2xhM2e had sedimented to the junction of 15%/45%.

Figure S4

The geometric mean titer of anti-hM2e IgG obtained from mice immunized by AH3-sfGFP-2xhM2e or LYRRLE-sfGFP-2xhM2e fusion protein once. The control group was the mice injected with PBS only. (N=5)
Supplementary materials

Figure S1

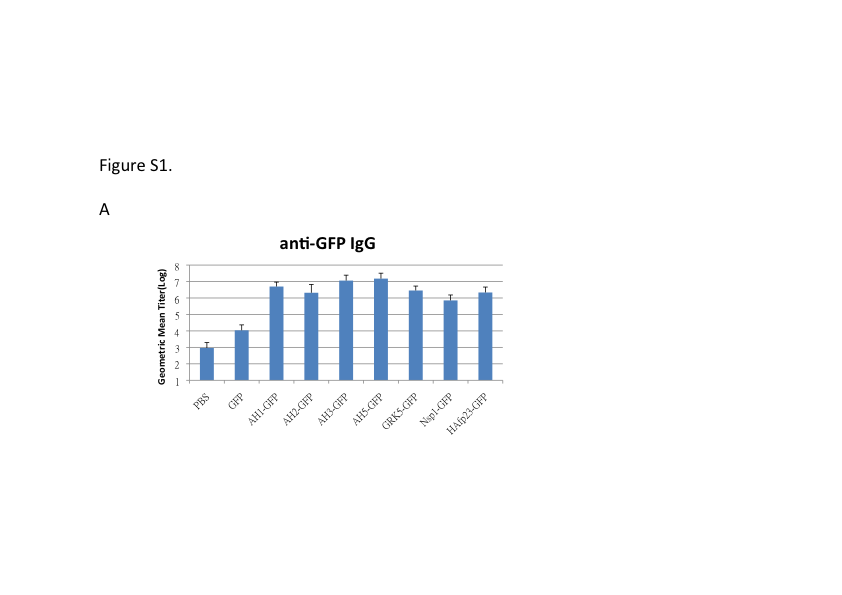

Figure S2

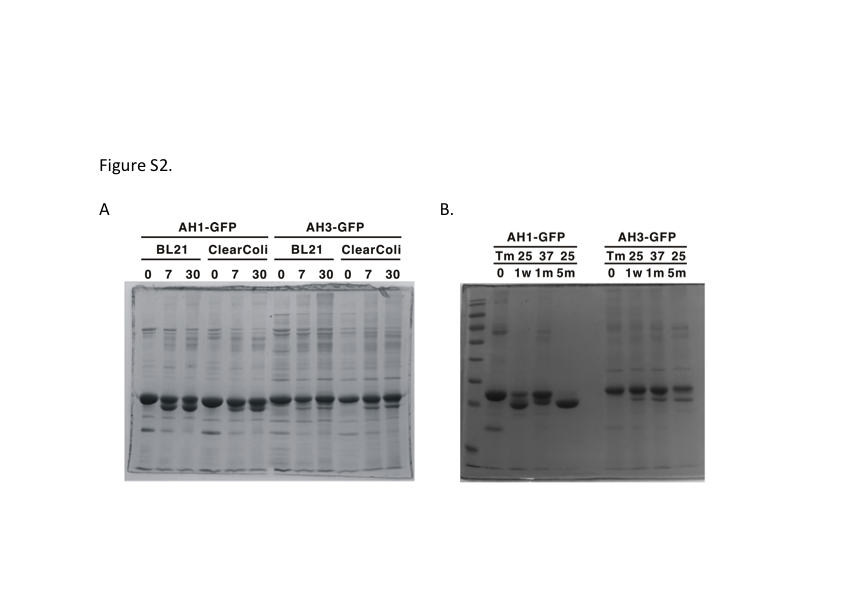

Figure S3

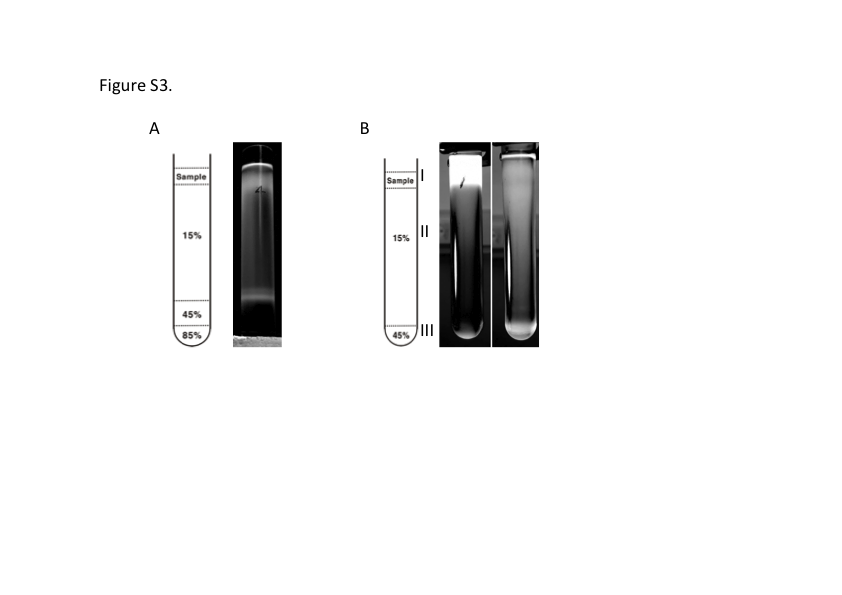

**Figure S4**

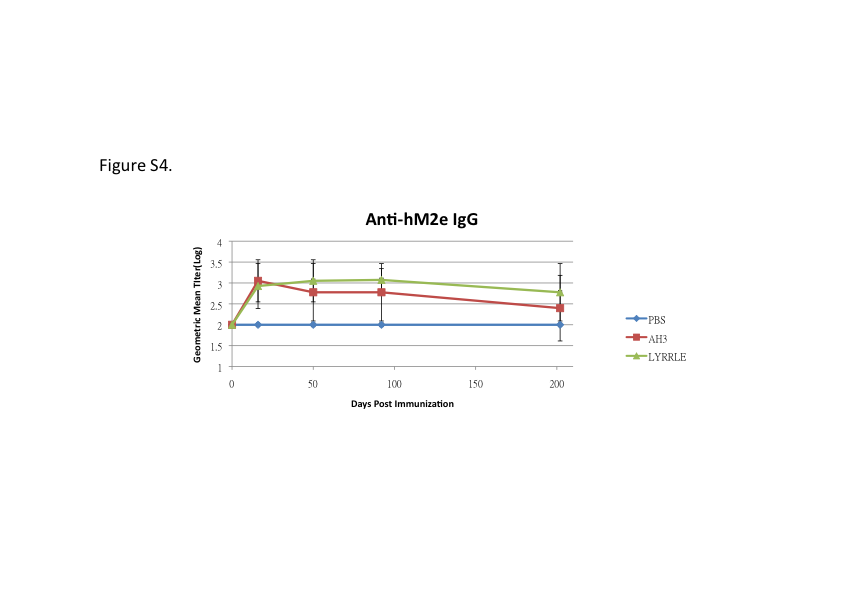

**Materials and Methods**

**Peptide information, expression and purification of recombinant protein**

The AH3 peptide sequence is DRLFFKCIYRRLKYGLKRG. The sequence of peptide for anti-hM2e antibody titer ELISA is SLLTEVETPIRNEWGSRSNGSSDC. The peptide inserted in the inertion site of sfGFP is SSLLTEVETPIRNEWGSRSN- GSSDSSGGSLLTEVETPIRNEWGSRSNGSSD. The protein expression vectors encoding target proteins were transformed into E coli competent cells using a heat shock transformation. Colonies of transformed bacteria with the desired vector were scraped from plate and inoculated in LB culture with an antibiotic. The bacterial culture was then growing continuously to OD600 between 0.5~0.7 before cooling down on an ice bath and protein expression was induced by 1mM IPTG at 20 ^o^C for 14-16 hours with 250 rpm shaking. After protein induction, bacteria were harvested by 5000 rpm centrifugation in a Sorvall SLC3000 rotor for 10 minutes. Bacterial pellet from 400 ml LB culture was re-suspended in 40 ml lysis buffer contains 10 mM Imidazole in 1XGF buffer (20 mM Na(PO_4_) pH7.4 and 300mM NaCl) for sonication. For Ni-NTA resin purification: 10 mM Imidazol in 1XGF buffer for bacteria lysis (Lysis buffer), 20 mM Imidazol in 1XGF buffer for column wash (Wash buffer), 500 mM Imidazol in 1XGF buffer for protein elution (Elution buffer). Bacteria were lysed using an ultrasonic sonicator (Misonix 3000) at 10 second on/20 second off cycles for 5 minutes in icy water. Insoluble debris was removed by centrifugation in 10000 rpm for 10 minutes using a Sorval SS34 rotor at 4 ^o^C. Soluble fraction containing the target protein was then used for purification by Ni-NTA resin as described in the user manual. Fusion protein eluted from Ni-NTA resin was then dialyzed against 1/2x GF buffer to remove excessive Imidazol and salt before immunization.

**Analysis of protein oligomerization and hydrophobicity**

Oligomerization state of AH3-GFP is first analyzed by protein concentration tube Vivaspin 2 with MWCO of 100, 300, or 1000 kDa from Sartorius. Protein samples were centrifuged in Vivaspin 2 at 1000g for 20 minutes, filtrate was analyzed by SDS-PAGE and then Coomassie blue staining for comparison. TEM images of AH3-GFP protein complex was obtained after the negative staining with phosphotungstic acid. Images were taken using Tecnai G2 Spirit Twin. For AH3-sfGFP-2xhM2e, the purified fusion protein was first crosslinked with Sulfo-SMCC(sulfosuccinimidyl 4-[N-maleimidomethyl]cyclohexane-1-carboxylate) at 4 μg/ml for 30 minutes before negative staining protocol. A sucrose density gradient was used to analyze VADEX oligomerization status. In a 13-ml polypropylene tube (Beckman cat#14287), the bottom was layered with 1 ml 85% (w/v) sucrose solution and topped with 2 ml 45% (w/v) sucrose solution and then 7 ml 15% (w/v) sucrose solution. The bacterial lysate for analysis was the soluble fraction described in the previous paragraph. Sudan III stock solution was prepared as 0.5% in isopropanol. Staining of bacterial membrane was by adding Sudan III stock solution into bacterial suspension before sonication at 1:100 dilution. One milliliter of the soluble fraction was layered on top of sucrose solutions. Protein samples were centrifuged at 35,000 rpm for 2 hrs in an SW41Ti swing bucket rotor using a Beckman ultracentrifuge, Optima L-90K. The centrifuge tubes were photographed in front of dark background in the present of a light bulb emitting 450 nm LED light. Images were processed in ImageJ from NIH by first splitting the image into R/G/B channels, then the green channel image were used for scanning all six samples from top to bottom using Plot Profile.

**Animal immunization and antibody titer determination**

The animal protocols had been approved by IACUC of Fu Ren Catholic University and mice were housed in the experimental animal center of Fu Ren Catholic University following SPF standards. Mice used in immunization procedures were between 8 to 9 weeks age. Mice were immunized through an intramuscular injection of purified protein preparation in 1 mg/ml concentration. For blood collection, mice were bled from a facial vein after pricked by the lancet. Sera collected were stored in -80 ^o^C before used in an ELISA assay. For each immunization, 20 μg purified protein was injected intramuscularly in the thigh of the hind limb. Blood was bled 14 days post immunization or planned dates for analysis using ELISA. For ELISA, the antigen was diluted with coating buffer for coating on high binding ELISA plate (Greiner Bio-One MICROLON high binding ELISA plate) , antigen concentration of hM2e peptide was 2.5 μg/ml and sfGFP was 1 μg/ml. The ELISA plate was incubated at 4 ^o^C overnight and then washed and blocked with 200 μl of blocking buffer containing 1% BSA in washing buffer. Washing buffer is composed by 10mM Na(PO_4_), 150mM NaCl at pH 7.4 with 0.05% Tween 20. Antiserum was diluted started from 1:100 and followed by sequential 4 folds dilution. Secondary antibody, Goat anti-mouse IgG with HRP conjugation was diluted at 1:5000 in blocking buffer. Chromogenic development using TMB was carried out following a standard protocol. Antibody titer was determined as the reciprocal value of the highest dilution that gives a reading of 0.1 above background in OD450.

**Protein structure modeling and modification**

Protein structure models of AH3 peptide monomer, dimer and tetramer were created using Ascalaph designer version 1.8.79. The intermolecular interaction forces between either monomers or dimers were calculated using the intermolecular energy command of Ascalaph designer. Side chain mutagenesis and interaction were modeled by Deep View/Swiss PdbViewer version 4.1.1. The solvent accessible surface area of the AH3 dimer was calculated using Jmol with a radius of 1.2 angstroms.
